# Optimisation of single-nuclei isolation and RNA sequencing of parasitic nematodes

**DOI:** 10.1101/2024.11.13.623282

**Authors:** Sarah K Buddenborg, Alison A Morrison, Anna C Bewsher, Manuela R Kieninger, Bee Ling Ng, Kirsty Maitland, Emily Hart, Dave J Bartley, Maria A Duque-Correa, Stephen R Doyle

## Abstract

Single-cell/nuclei transcriptomics has revolutionised our understanding of cell, tissue, and organismal biology in humans and model organisms across health and disease. However, for non-model species, reassessment and refinement of conventional methods is needed to address novel technical and biological challenges. Here, we evaluate three established nuclei isolation protocols and, building on these experiences, develop an optimised protocol for single-nuclei isolation of parasitic nematodes (SNIP). SNIP recovered 1.7- to 4.1-fold more nuclei than the three established protocols, and 3.2-fold more RNA per nucleus than the best-performing of these. We validated SNIP across macroscopic and microscopic life stages and both sexes of two phylogenetically distinct gastrointestinal nematodes, *Haemonchus contortus* and *Trichuris muris*, and used it to generate single-nuclei RNA-seq data revealing sex- and stage-specific gene expression and putative tissue-specific expression in both species. By enabling flexibility with frozen samples and consistent recovery across diverse parasitic nematode life stages, SNIP provides a practical foundation for building cell and nuclei atlases in other non-model species, toward advancing understanding from fundamental biology to infection and disease.

## Introduction

With the increasing affordability and availability of high-throughput sequencing and commercial single-cell (sc) and single-nuclei (sn) transcriptomic platforms, cellular-resolution approaches to investigate the biology of non-model organisms are becoming more accessible. Applying sc/snRNA-seq to non-model organisms, however, introduces biological and technical challenges that may need to be overcome. The biological richness of non-model species brings a diversity of cell and tissue types that vary in morphology and composition and differ in their sensitivities to cellular and nuclear dissociation; cell size, viability, dissociation feasibility and the presence of rigid cell walls or other protective structures must each be assessed per species (Woo and Eyun, 2025). Sample access may also be constrained, whether by geography, seasonality, or the inability to culture an organism outside its natural setting, making reliance on preserved rather than fresh material a practical necessity rather than a convenience. Addressing these challenges requires organism-specific biological knowledge, and for most species no established protocol exists to draw on. How well methods developed for model organisms transfer to this diversity remains largely untested.

A field with exclusively non-model organisms that stands to benefit greatly from understanding biology at a single-cell resolution is parasitology. scRNA-seq of single-cell eukaryotic parasites such as *Plasmodium spp.* and *Trypanosoma spp.* has been particularly successful, with major advances in understanding life-stage-specific gene expression patterns and transitions during their life cycles, mechanisms of infection and disease progression, and the identification of new drug targets to interrupt transmission (Briggs et al., 2023, 2021; Howick et al., 2022, 2019; Real et al., 2021; Reid et al., 2018; Vigneron et al., 2020). In contrast, the application of scRNA-seq to multicellular parasites has encountered distinct challenges due to vast differences in life-history traits and hosts, and to the great diversity in body plan and cellular composition between and even within species throughout their life cycles. Multicellular parasites are exposed to variable environmental conditions inside and outside the host, necessitating a tough outer coating (i.e., cuticle, tegument, exoskeleton), which presents a major obstacle to single-cell/nuclei isolation. Further, because most species cannot be cultured *in vitro*, access to obligate parasitic stages can be geographically restricted and difficult or impossible to collect without sampling post-mortem. Despite these challenges, there are a growing number of single-cell life-stage atlases for parasitic helminths, including the trematodes *Schistosoma mansoni* (Attenborough et al., 2024; Diaz Soria et al., 2024, 2020; Li et al., 2021) and *Fasciola hepatica* (Puckelwaldt et al., 2024), and the cestode *Echinococcus multilocularis* (Loos et al., 2026). Similarly, cell atlases have been generated for *Brugia malayi* microfilariae (Henthorn et al., 2023), *Ascaris suum* intestinal tissue (Tyagi et al., 2024), *Haemonchus contortus* embryos (Korhonen et al., 2024), and *Heligmosomoides bakeri* mixed-sex adults (Pollo et al., 2024). These studies often utilise dissociation protocols developed for mammalian tissues, although it is unclear how transferable and efficient these methods are across diverse species and life-cycle stages of parasitic helminths.

In this study, we tested and compared three widely used single-nuclei isolation protocols for snRNA-seq in parasitic nematodes. We focused on isolating single nuclei rather than single cells because frozen material can be used more readily, providing several advantages, including the ability to sample across life stages, collect and store samples over time, and process them simultaneously to minimise batch effects. Building on our experiences from the three-protocol comparison, we have developed a new nuclei isolation protocol and tested it across multiple life-cycle stages and sexes of two gastrointestinal parasitic nematodes, *H. contortus* and *Trichuris muris*. We assessed the nuclei preparations from our protocol using the Parse Biosciences Evercode WT Mini platform for snRNA-seq and Illumina sequencing. Here, we provide the protocol and discuss some of the challenges and opportunities of sc/snRNA-seq approaches to advance the adoption of this technology in non-model organisms and its applications to better understand helminth biology.

## Materials and Methods

### Trichuris muris

#### Mice husbandry and ethical approval

NSG (NOD.Cg-*Prkdc^scid^ Il2rg^tm1Wjl^*/SzJ) mice were maintained under specific pathogen-free conditions, under a 12 h light/dark cycle at a temperature of 19-24°C and humidity between 40 and 65%. Mice were naive before the studies described here. Experiments were performed under the regulation of the UK Animals Scientific Procedures Act 1986 under the Project license P77E8A062 and were approved by the Wellcome Sanger Institute Animal Welfare and Ethical Review Body.

#### T. muris infection of mice

Infection and maintenance of *T. muris* were conducted as described (Duque-Correa et al., 2019; Wakelin, 1967). Briefly, NSG mice (6-8 weeks old) were orally infected under anaesthesia with isoflurane with a high dose of embryonated eggs (n = 400) from *T. muris* E-isolate.

#### Adult worm collection

Thirty-five days post-infection, mice were culled by cervical dislocation and the caecum and proximal colon were removed. The caecum was split and washed in RPMI-1640 supplemented with 500 U/mL penicillin and 500 μg/mL streptomycin (all from Gibco, UK). Worms were removed using fine forceps, and males and females were sorted into separate 1.5 mL nuclease-free microcentrifuge tubes. All liquid was removed from the worms in the tubes, and the tubes were snap frozen in liquid nitrogen and stored at -80°C.

### Haemonchus contortus

#### Sheep husbandry and ethical approval

Sheep used for experimental infections of *H. contortus* were born and raised under worm-free conditions at the Moredun Research Institute, UK. Sheep were humanely euthanised under Schedule 1 of the Animals (Scientific Procedures) Act. All experimental procedures were examined and approved by the Moredun Research Institute Animal Welfare and Ethics Review Board and were conducted under approved UK Home Office licences (PPL 60/03899, experimental code identifier E46/11).

#### Infection, culturing, and collection of L3 larvae

Sheep were infected orally with 5,000 *H. contortus* MHco3(ISE) L3. Faeces were collected 21 days post-infection and coprocultured for 14 days to allow development from egg to L3, after which the larvae were recovered using the Baermann technique and stored at approximately 8°C in tap water. Larvae were quantified by mixing the 50 mL stock thoroughly, diluting a 1 mL subsample to 10 mL in a volumetric flask, and counting twelve 10 µL streaks. The highest and lowest counts were discarded and the mean of the remaining ten used to calculate the total: mean larvae per 10 µL × 100 gives larvae per mL, × 10 gives the count in the 10 mL dilution (equivalent to 1 mL of the original stock), × 50 gives the total in the 50 mL stock.

To collect the larvae, water containing the larvae was vacuum-filtered through a 10 µm cell strainer (pluriSelect; cat. 43-50010-03) fitted to a 250 mL filter flask and washed with 50 mL of 1x M9 buffer. Larvae were scraped off the cell strainer using a micro spatula (Fisherbrand; cat. 11523482), put in a 1.5 mL nuclease-free microcentrifuge tube, immediately submerged in liquid nitrogen, and stored at -80°C.

#### Adult worm collection

Adult worms were collected from the abomasum of sheep at necropsy 28 days post-infection. Worms were rinsed in physiological saline (0.85% NaCl) and sexed based on morphology, before male and female adults were sorted into separate 1.5 mL nuclease-free microcentrifuge tubes using a needle. The tubes were submerged in liquid nitrogen and stored at -80°C.

### Established nuclei isolation protocols evaluated in this study

#### Overview of protocol evaluation

Four nuclei isolation protocols were compared: three established protocols (Celegans, HCA and Frankenstein) and the optimised protocol developed here (SNIP). Each was performed in triplicate (biological replicates, n = 3), using ten adult female *H. contortus* per replicate as a standardised input across all four protocols. All four protocols were performed by the same operator, using the same batch of worms, to minimise variation unrelated to the protocols themselves.

Unless otherwise stated, all reagents and equipment were pre-chilled, samples were kept on wet ice throughout, and all centrifugation steps were performed at 4°C. The three established protocols were carried out as published, except where deviations are stated explicitly in the sections below.

Following selection of the optimised protocol, SNIP was validated across additional species and life stages: 10 adult female, 20 adult male and approximately 250,000 L3 *H. contortus*, and 25 adult female and 25 adult male *T. muris*, each performed in triplicate. Inputs for these samples were matched by approximate volume per tube rather than by worm number, owing to the large differences in body size between stages and species.

#### Frankenstein protocol

Nuclei were isolated using the Frankenstein protocol (Martelotto, 2020), which combines strategies from five protocols developed for nuclei isolation from mammalian brain tissue (Habib et al., 2017, 2016; Hu et al., 2017; Lacar et al., 2016; Lake et al., 2016). Worms were minced with a razor blade on wet ice and transferred to a 1.5 mL tube containing 500 µL chilled Nuclei EZ Lysis Buffer (Sigma-Aldrich; cat. NUC101-1KT). The sample was homogenised with 15 strokes of a plastic micro-pestle (Fisherbrand; cat. FS7495210590), a further 1 mL of chilled EZ Lysis Buffer was added, and the homogenate was incubated on ice for 5 min with gentle mixing using a wide-bore pipette tip during the incubation. The homogenate was filtered through a 70 µm strainer, and nuclei were pelleted at 500 × *g* for 5 min at 4 °C, resuspended in 1.5 mL EZ Lysis Buffer, incubated on ice for 5 min, and pelleted again under the same conditions. Nuclei were resuspended in 1% BSA-PBS containing 0.2 U/µL SUPERase In^TM^ RNase inhibitor (Thermo Fisher; cat. AM2694), filtered through a 35 µm cell strainer, and sorted by FACS.

#### Celegans protocol

The Celegans protocol was developed for the free-living nematode *C. elegans* by Truong *et al*. (2023) and described in detail by Levenson *et al*. (2023). The protocol specifies a compact 30 µL worm pellet as input. Worms were transferred to a pre-chilled 1.5 mL glass Dounce tissue grinder and homogenised in 450 μL ice-cold modified FA cell lysis buffer (50 mM HEPES/NaOH pH 7.5, 1 mM EDTA, 0.1% Triton X-100, 150 mM NaCl, with 0.5× protease inhibitor [Roche; cat. 11697498001] and 0.2 U/µL RNaseOUT^TM^ Recombinant Ribonuclease Inhibitor [Thermo Fisher; cat. 10777019] added fresh) and homogenised with ten strokes of the tight pestle. The homogenate was centrifuged at 100 × *g* for 1 min at 4 °C and the nuclei-containing supernatant retained. A further 300 µL modified FA buffer was added to the residual debris, which was homogenised with ten strokes of an Eppendorf Dounce homogeniser using a corkscrew motion and clarified as above; this was repeated once, giving 30 strokes in total. Pooled supernatants were centrifuged, and nuclei were washed in 1% BSA-PBS and filtered through a 40 µm Flowmi tip strainer (Sigma Aldrich; cat. BAH136800040) before sorting by FACS.

#### Human Cell Atlas (HCA) protocol

Nuclei were isolated using the protocol of the Human Cell Atlas team at the Wellcome Sanger Institute (Stephenson, E., Prigmore, E., Oszlanczi, A., Zhou, D., 2023), adapted from Nadelmann *et al*. (2021). Unlike the other protocols assessed, tissue was not minced or chopped before homogenisation. Buffers were prepared as described in the source protocol: nuclei isolation buffer 1 (NIM1; 250 mM sucrose, 25 mM KCl, 5 mM MgCl, 10 mM Tris in nuclease-free water) was stored at 4 °C, and homogenisation buffer (HB) was prepared fresh on ice on the day of use by supplementing NIM1 with 1 µM DTT, 0.4 U/µL RNasin Plus (Promega; cat. N2611), 0.2 U/µL SUPERase·In RNase inhibitor (ThermoFisher; cat. AM2694) and 0.1% Triton X-100. The 1× protease inhibitor specified in the source protocol was omitted, based on advice from Sanger core single-cell team. Worms were transferred directly into a Wheaton Dounce tissue grinder (DWK Life Sciences; 357538) containing 1 mL HB and homogenised on ice with up to 20 strokes of the loose (A) pestle until no resistance was felt, the pestle was rinsed with 500 µL HB, and the process was repeated with the tight (B) pestle. The homogenate was passed through a 40 µm cell strainer (Thermo Fisher, cat. 087711), the homogeniser rinsed with a further 500 µL HB, and nuclei were pelleted at 500 × *g* for 6 min at 4 °C and washed in 500 µL wash buffer (1× PBS, 5% BSA, 40 U/µL RNasin protector) for 2 min on ice. Debris was then removed by density gradient centrifugation (225 µL of nuclei suspension and 300 µL 90% Percoll-1x PBS were mixed by inverting) substituting Percoll (Sigma-Aldrich, cat. P4937) for the Lymphoprep specified in the source protocol, at 20,000 × *g* for 15 min at 4 °C.

#### SNIP: an optimised protocol for single-nuclei isolation from parasitic nematodes

Tissue disruption was performed differently for macroscopic and microscopic life stages, with samples kept frozen throughout in both cases. For adult worms (*H. contortus* and *T. muris*), frozen worms were cut into pieces <2 mm directly in the sample tube on dry ice using spring scissors pre-chilled in a 50 mL Falcon tube on dry ice. For microscopic stages (*H. contortus* L3), approximately 250,000 larvae were ground in a porcelain mortar and pestle (Fisher Scientific; cat. 12383338) pre-cooled on dry ice. Disrupted tissue was transferred to a 1 mL Dounce tissue grinder containing 700 µL Nuclei EZ Lysis Buffer (Sigma-Aldrich; cat. NUC101) with the addition of 25 µg/mL DNase I (STEMCELL Technologies; cat. 07900), 5 mM MgCl_2_, and 0.2 U/µL SUPERase·In RNase inhibitor using a glass Dounce tissue grinder. The loose Dounce tissue grinder pestle was applied five times, followed by three strokes of the tight Dounce tissue grinder pestle. The homogenate was held on ice for up to 4 min, or until large undissociated tissue and debris had settled, and the nuclei-containing supernatant was transferred to a 5 mL Protein LoBind tube (Eppendorf). Lysis buffer was replenished to the original volume, three further strokes of the tight pestle were applied, and the supernatant was collected again after settling. Homogenisation with the tight Dounce tissue grinder pestle was repeated up to three more times or until the settled homogenate was clear. Pooled nuclei were pelleted at 500 × *g* for 5 min at 4 °C, washed once with the 2% BSA-PBS wash buffer containing 0.2 U/µL SUPERase·In and 5 mM EDTA, and then filtered through a 30 µm CellTrics filter (Sysmex; cat. 04-004-2326). The filter was washed again with wash buffer to maximise the nuclei recovery. Nuclei were then sorted using FACS to remove small debris.

#### Nuclei staining, flow cytometry and FACS gating

Nuclei were assessed and sorted on a Bigfoot Spectral Cell Sorter (ThermoFisher). For each sample, a 50 µL aliquot of nuclei was retained unstained as a negative control for setting fluorescence gates. The remaining suspension (∼500 µL) was stained with 5 µL 10,000× SYBR Green I (ThermoFisher; cat. S7567). Fluorescence was excited with a 488 nm laser and detected through a 507/19 filter. Nuclei were gated on SYBR Green I signal alone, without scatter-based gates, to avoid excluding the unusually small and large nuclei observed by microscopy. Sorted nuclei were collected into pre-chilled 5 mL Protein LoBind tubes (Eppendorf) blocked with 1% BSA and containing 2% BSA-PBS wash buffer (composition as above), and kept on ice.

#### Post-sort nuclei fixation and cryostorage (Evercode Nuclei Fixation kit)

After sorting, the nuclei were stained with trypan blue and the concentration was determined using a C-Chip disposable haemocytometer (NanoEntek; cat. DHC-N01) and then pelleted at 500 x *g* for 10 min at 4 °C. Nuclei were fixed using the Evercode Nuclei Fixation kit (Parse Biosciences) according to user manual v2.0.2 (Rebboah and Biosciences, 2023), with three modifications. First, the 15 mL polypropylene tubes specified in the protocol were replaced with 5 mL Protein LoBind tubes, blocked with 1% BSA in nuclease-free water (BSA-NFW); tubes were filled with 1% BSA-NFW, incubated for 30 min at room temperature, and dried in a laminar flow cabinet. All centrifugation steps were performed at 500 × *g*. Second, the two 40 µm straining steps were omitted to maximise nuclei retention, as our nuclei concentration was at the lower limit of the recommended input range (100,000 nuclei), and no nuclear clumping was observed by microscopy. Third, nuclei were resuspended in 50 µL rather than 150 µL Nuclei Buffer, with the DMSO volume adjusted proportionally (5% v/v). Fixed nuclei were frozen in a Corning CoolCell container and stored at −80 °C, until all preparations had been collected and could be processed together.

#### snRNA-seq library preparation and Illumina sequencing

Combinatorial barcoding, cDNA amplification and library preparation were performed using the Evercode WT Mini v2 kit (Parse Biosciences) according to user manual v2.0.1. A 10 µL aliquot of fixed nuclei from each sample was thawed at 37 °C and nuclei were counted manually using a C-Chip haemocytometer. Nuclei concentrations were entered into the Parse Biosciences Evercode WT Mini Sample Loading Table v1.2.0, which calculates the volume of sample stock required from the number of samples, the target number of barcoded nuclei, and the proportion of the library contributed by each sample. After three rounds of barcoding, nuclei were pooled, counted manually using a C-Chip haemocytometer, and divided into two equal sub-libraries. Sub-library nuclei were lysed and a fourth, sub-library-specific barcode was added by PCR extension, which also adds Illumina-compatible adapters. Seven cycles were used in the second cDNA amplification step and 13 or 10 cycles in the sub-library index PCR, as specified by the kit for the nuclei numbers used for experiments 1 and 2, respectively. cDNA sub-libraries and final libraries were assessed using the HS D5000 TapeStation assay (Agilent) and quantified using the HS dsDNA Qubit assay (ThermoFisher).

The resulting libraries were combined with 5% PhiX, and then each was sequenced on an Illumina NovaSeq 6000 lane using 150 bp paired-end reads, generating 316,638,084 and 307,516,986 reads for Library 1 and Library 2, respectively.

#### snRNA-seq data processing, clustering and cluster annotation

Raw sequencing data were first assessed using FastQC (v0.12.1) (https://www.bioinformatics.babraham.ac.uk/projects/fastqc/) and MultiQC (v1.17) (Ewels et al., 2016) before analysis using the Parse Biosciences split-pipe (v1.9.0) pipeline. Reference genomes and annotations were downloaded from WormBase ParaSite release WBPS19 for *H. contortus* (PRJEB506; Doyle et al., 2020) and *T. muris* (PRJEB126; Foth et al., 2014). Reference indices were built for each species separately using *split-pipe --mode mkref* (v1.9.0, Parse Biosciences) from the genome (fasta) and annotation (gff3) files. Because nuclei from both species were barcoded together within each experiment, reads from each library were mapped separately against each reference using *split-pipe --mode all*, with *post_min_map_frac* set to 0.01 as recommended for species-separate experiments and *pre_check_name_match* set to False to accommodate Sanger-generated FASTQ headers. For experiment 2, the two sub-libraries were processed individually and combined using *split-pipe --mode comb*. Mapping statistics, nuclei identified, and genes and transcripts per nucleus were taken from the resulting summary reports.

Parse Biosciences split-pipe output files were uploaded into Trailmaker^TM^ (https://app.trailmaker.parsebiosciences.com/; pipeline v1.5.0, Parse Biosciences) to use the “Cell size distribution filter”. This filter uses a barcode rank plot to determine the minimum number of transcripts per cell for each sample. Filtering and subsequent analysis, including generation of UMAP plots, and identification of nuclei cluster marker genes were performed using Seurat (v5.0.0) (Hao et al., 2024) in R (v4.4.0). Nuclei were filtered using the cell size distribution cut-off and then doublets were removed after running scDblFinder (Germain et al., 2021). Raw counts were normalised with Seurat’s *LogNormalize* method and the top 2000 highly variable genes (HVGs) were selected using the *vst* method. Then, the data was scaled using the *ScaleData()* function and a Principal Component Analysis (PCA) was run using 50 principal components. Harmony (Korsunsky et al., 2019) integration within seurat was selected to correct for batch effects and this corrected data was used for generating UMAPs. The *FindNeighbors* function computed k-nearest neighbors (KNN) and shared nearest neighbor (SNN) graphs before running *FindClusters* with the resolution set to 0.8. UMAPs were visualised, combining all life stages per species and subsequently coloured by life stage or cluster ID. To infer the biological relevance or putative tissue type of each cluster, marker genes were obtained using Seurat’s *FindAllMarkers* function. Gene descriptions and putative function for the marker genes were cross referenced with *Caenorhabditis elegans* orthologs identified using WormBase ParaSite Biomart (Howe et al., 2017).

#### Statistical analysis

All statistical analyses were performed in R (v4.5.0). Nuclei yields are reported as the mean of three biological replicates ± standard deviation. Pairwise comparisons of nuclei yield between protocols, and of nuclei per worm between species, were made using two-tailed Welch’s *t*-tests, which do not assume equal variances between groups. Given the small number of replicates (n = 3 per group), normality could not be meaningfully assessed and these tests are reported as indicative rather than confirmatory.

Nuclei density in FACS data was analysed in R using flowCore (v2.22.1). Events above the SYBR Green I gate were log -transformed and peak apexes identified from a kernel density estimate. Each peak was fitted by maximum likelihood on unbinned events in a window around its apex, modelling the density as a truncated normal plus a uniform baseline; window width and baseline slope were selected per peak by fit convergence. Fold changes are ratios of fitted peak means, with 95% percentile bootstrap intervals from 500 resamples of events. Sensitivity to window width was assessed by refitting at 0.8× and 1.25× the chosen width.

## Results

### Comparison of three existing nuclei isolation protocols

To optimise a protocol for generating single nuclei suspensions from parasitic nematodes, we first compared three established isolation protocols: (i) a *C. elegans* nuclei isolation protocol (hereinafter “Celegans”) (Levenson et al., 2023; Truong et al., 2023), (ii) a Human Cell Atlas (HCA) protocol (Stephenson, E., Prigmore, E., Oszlanczi, A., Zhou, D., 2023), and (iii) the Frankenstein protocol (Martelotto, 2020). An overview of the four key steps of the three protocols is shown in **Fig. 1A**, which includes tissue disruption, cell lysis and dissociation, homogenisation, and debris removal. Each protocol was performed using ten frozen adult female *H. contortus* worms to enable direct comparison.

**Fig. 1.**
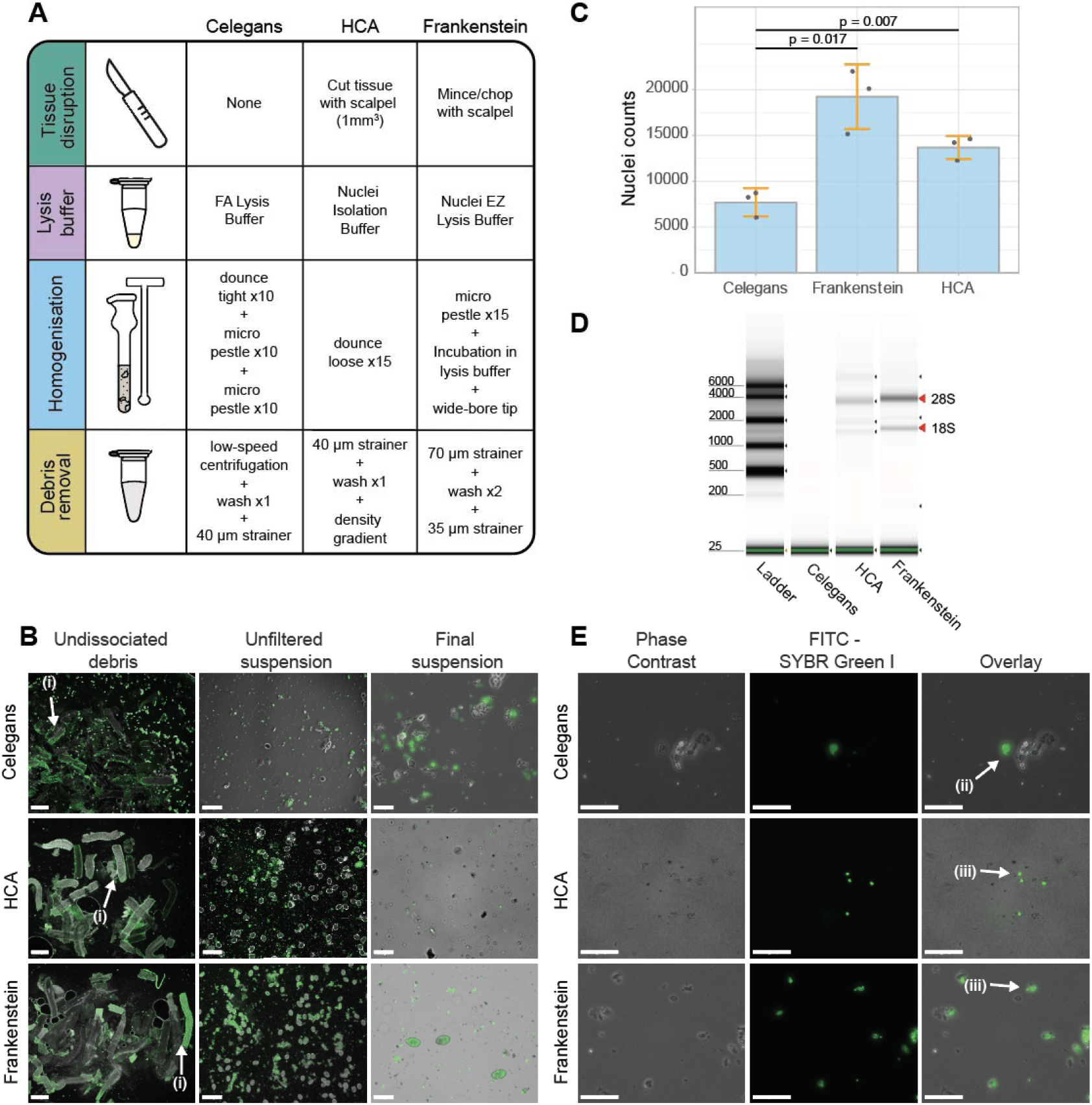
Isolation of nuclei from ten adult female *Haemonchus contortus* worms using three existing protocols. A) Schematic comparison of the Celegans, HCA and Frankenstein nuclei isolation methods. B) Representative images from each protocol of the undissociated debris remaining after homogenisation, the unfiltered suspension, and the final suspension. Arrows (i) indicate empty cuticle fragments remaining after homogenisation. Images are overlays of phase-contrast and fluorescence microscopy of SYBR Green I-stained samples; scale bars = 500, 250 and 50 µm for the undissociated debris, unfiltered suspension and final suspension columns, respectively. C) Manual nuclei counts. Bars show the mean of three replicates, error bars the standard deviation, and points the individual replicates. *p*-values are from two-tailed Welch’s *t*-tests; HCA and Frankenstein did not differ significantly (*p* = 0.099). D) Representative Agilent HS RNA TapeStation gel of RNA extracted from the nuclei suspension of each protocol. E) Representative phase-contrast, SYBR Green I fluorescence, and merged images of the final nuclei suspensions. Scale bars = 50 µm. Arrow (ii) indicates a broken nuclear membrane leaking stained nucleic acid (Celegans); arrows (iii) indicate nuclei of differing size (Frankenstein).

#### Tissue disruption and lysis

Tissue disruption is particularly important for nematodes, whose highly cross-linked acellular cuticle protects the animal from its environment. The HCA and Frankenstein protocols mince tissue with a scalpel or razor blade before lysis, whereas the Celegans protocol includes no disruption step, relying instead on homogenisation to expose cellular contents — an approach suited to the small size of *C. elegans* (∼1 mm × 50 µm) but untested on larger parasitic species. Each protocol uses a different lysis buffer, though all contain a detergent to disrupt membranes, a pH stabiliser, salts to provide ionic strength, and agents that protect RNA from degradation. Notably, the commercial lysis buffer used in the Frankenstein protocol is not supplemented with an RNase inhibitor, which is added only at the wash and resuspension stage. Because lysis acts in concert with mechanical homogenisation, its contribution cannot readily be assessed in isolation.

#### Homogenisation efficiency varies markedly between protocols

Examining samples microscopically after homogenisation revealed clear differences in the completeness of tissue disruption (**Fig. 1B**). Empty cuticle debris predominated in the Celegans protocol, indicating that homogenisation had gone to completion, consistent with its use of ten strokes of the tight Dounce pestle followed by two sets of ten strokes with a plastic micro-pestle, with suspended nuclei collected between sets to avoid over-homogenisation. In contrast, large worm fragments remained prominent in both the HCA and Frankenstein homogenates, indicating incomplete homogenisation despite the HCA protocol’s two rounds of up to 20 Dounce strokes with the loose and tight pestles, and the Frankenstein protocol’s reliance on a plastic micro-pestle with wide-bore mixing and two 5-minute lysis incubations.

#### Large debris was removed effectively, but small debris persisted in all three protocols

Homogenisation generates substantial debris, including acellular material and the products of cell lysis. Large debris (>30 µm) can cause clogs or wetting failures in microfluidics-based platforms (*Best Practices to Minimize Chromium Next GEM Chip Clogs and Wetting Failures*, 2022), though it is less problematic for combinatorial indexing approaches in which nuclei are distributed across microtitre plates. Nonetheless, small debris (<30 µm) may contain ambient RNA, increasing transcriptional background noise (Madissoon et al., 2019) and should therefore be removed. All three protocols removed large debris effectively using cell strainers. Small debris persisted in all three and was most abundant in the Celegans protocol (**Fig. 1B**). The density gradient step in the HCA protocol — for which we substituted Percoll for the Lymphoprep specified in the published protocol — removed no more small debris than the simpler wash steps used by the other protocols, while doubling the total protocol time (**Fig. 1B**).

#### Nuclei yield, RNA recovery and morphology

Nuclei suspensions were assessed on total nuclei recovered, RNA yield, and nuclei morphology. From ten adult female *H. contortus*, the Celegans, HCA and Frankenstein protocols recovered a mean of 7,700 ± 1,540, 19,227 ± 3,520 and 13,678 ± 1,265 nuclei (mean ± SD, n = 3), respectively (**Fig. 1C**). Yields differed significantly between Celegans and HCA (*t*(2.7) = 5.20, *p* = 0.017) and between Celegans and Frankenstein (*t*(3.9) = 5.20, *p* = 0.007), but not between HCA and Frankenstein (*t*(2.5) = 2.57, *p* = 0.099).

RNA extraction of nuclei preparations revealed that the Frankenstein protocol showed the most prominent 18S and 28S ribosomal RNA peaks on the High Sensitivity RNA TapeStation gel (**Fig. 1D**). The Frankenstein protocol recovered the most RNA (11,790 pg), followed by HCA (7,770 pg) and Celegans (5,940 pg). All concentrations fell below the threshold at which an RNA integrity number (RIN) can be reported. This reflects the low RNA input rather than degradation: RIN calculation depends on resolving the 18S and 28S ribosomal peaks against background, which is not possible at these concentrations. Visual inspection of the High Sensitivity RNA TapeStation traces nonetheless showed clear 18S and 28S peaks, most prominently for the Frankenstein sample (**Fig. 1D**), indicating acceptable integrity rather than degradation. Relative to the nuclei counts of the corresponding replicates, this equates to 0.52, 0.41 and 0.81 pg RNA per nucleus for Celegans, HCA and Frankenstein, respectively.

Nuclei morphology differed most clearly between protocols (**Fig. 1E**). Nuclei from the Celegans protocol showed notable degradation of the nuclear envelope, consistent with its lower RNA recovery; the arrow in the FITC-SYBR Green I and overlay panels (ii) indicates a broken nuclear membrane leaking fluorescently stained nucleic acid. Most nuclei from the Frankenstein protocol had smooth, intact membranes and spanned a range of sizes (iii), suggesting recovery from diverse cell types. Nuclei from the HCA protocol appeared largely intact, with smooth, well-defined membranes, but were predominantly small and uniform in size, with few of the larger nuclei recovered by the Frankenstein protocol.

In summary, the Frankenstein protocol generated the most RNA per nucleus and recovered morphologically robust nuclei spanning the widest range of sizes, suggesting capture from a more diverse set of cell types, with a nuclei yield that did not differ significantly from that of the HCA protocol.

### Optimisation of the Single Nuclei Isolation of Parasites (SNIP) protocol

Drawing on the limitations identified above, we developed an optimised protocol for parasitic nematodes (**Fig. 2A**; **Table 1**).

**Fig. 2.**
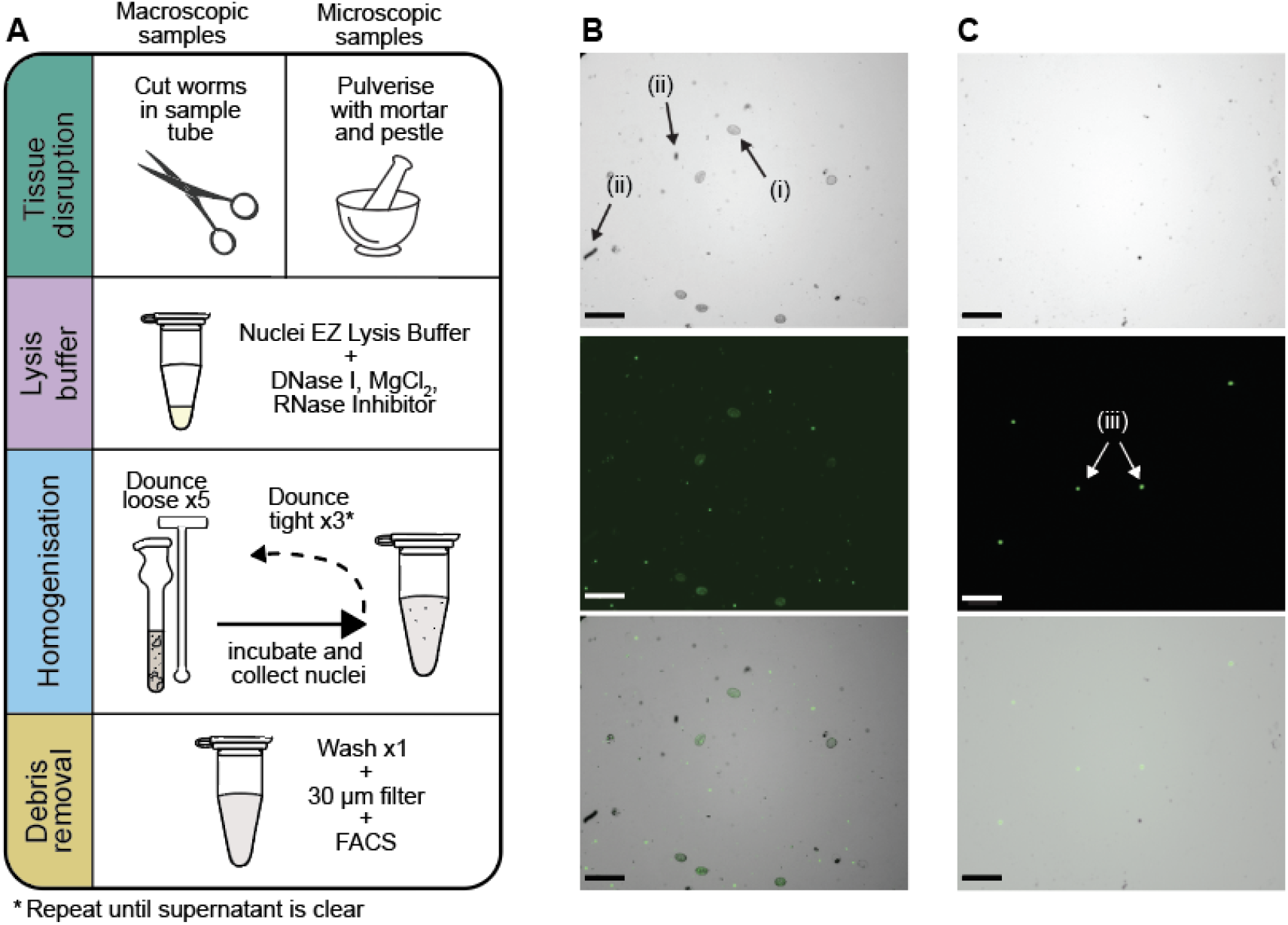
Overview of the Single Nuclei Isolation of Parasites (SNIP) protocol. A) Schematic of the optimised SNIP protocol, showing the alternative tissue disruption steps used for macroscopic (adult worms) and microscopic (larvae) samples; subsequent steps are common to both. B) Representative phase-contrast, SYBR Green I fluorescence, and merged images of an *H. contortus* adult female nuclei suspension after washing and filtering, before FACS. Arrows (i) indicate intact eggs and (ii) cuticle fragments remaining in the suspension. Scale bar = 100 µm; C) The same suspension after debris removal by FACS, shown as phase-contrast, SYBR Green I fluorescence, and merged images. Arrows (iii) show clean, fluorescent nuclei. Scale bars = 100 µm.

**Table 1.** Limitations of existing protocols and the corresponding modifications in SNIP.

| Protocol step | Previous limitation | Modification |
| --- | --- | --- |
| Tissue disruption | Mincing tissue on wet ice thaws worms immediately, risking RNA degradation and loss of nuclear quality; transferring tissue between a petri dish and the sample tube lost material. | Frozen worms were cut into <2 mm pieces directly in the sample tube on dry ice using pre-chilled spring scissors. Microscopic stages were ground in a dry-ice-cooled mortar and pestle. |
| Lysis | DNA released from lysed nuclei is "sticky" and causes intact nuclei to clump; in the Frankenstein protocol the commercial lysis buffer was not supplemented with an RNase inhibitor, which is added only at the wash stage. | Nuclei EZ Lysis Buffer* was supplemented with DNase I, MgCl<sub>2</sub> and RNase inhibitor. DNase I to digest free DNA, preventing clumping; MgCl<sub>2</sub> is required for DNase I activity and stabilises chromatin; adding RNase inhibitor at lysis protects RNA from the point of release rather than after two washes. |
| Homogenisation | The Frankenstein and HCA protocols left substantial undigested tissue, while homogenisation in the C elegans protocol was harsh enough to degrade nuclear membranes. | 1 mL Wheaton Dounce tissue grinder, five strokes with the loose pestle followed by three with the tight pestle, repeated up to three times. The loose pestle exposed tissue that scissors did not disrupt and released the first nuclei; the tight pestle lyses remaining tissue and additional nuclei. |
|  | Continuous homogenisation exposed already-released nuclei to repeated shear. | Each set of strokes was followed by incubation on ice, allowing large fragments to settle while nuclei in the supernatant are collected. This removed nuclei from further homogenisation and allowed lysis to continue within larger tissue fragments. |
| Debris removal | Each additional wash or filtration step lost nuclei and added to the processing time. | A single wash in 2% BSA-PBS with 0.2 U/μL RNase inhibitor and 5 mM EDTA. Pre-filtering through a 70 μm strainer is unnecessary because large debris remains settled in the Dounce tube. |
|  | Strainers alone did not sufficiently remove small debris from any of the three protocols, and the Percoll density gradient removed no more than a simple wash while doubling protocol time. | A 30 μm CellTrics filter followed by FACS. FACS was the only method that removed small debris effectively. Washing the filter through with additional buffer increased nuclei recovery. |
| Miscellaneous | Nuclei adhered to untreated plastic surfaces; fixed-angle rotors pelleted nuclei against the tube wall, making the pellet easy to disturb. | We recommend low-binding, BSA-blocked pipette tips and 5 mL Protein LoBind tubes, nuclei pelleted in a swing-bucket rotor results in a lower likelihood of disturbing the pellet during washes. |

#### Keeping tissue frozen during disruption

We disrupted tissue while frozen, using pre-chilled spring scissors directly in the sample tube for macroscopic stages and a dry-ice-cooled mortar and pestle for microscopic stages, avoiding the thawing that occurs when tissue is minced on wet ice (Table 1).

#### Preventing nuclei clumping and RNA degradation

We supplemented the lysis buffer with DNase I and MgCl, and added an RNase inhibitor to both lysis and wash buffers rather than to the wash buffer alone as in the Frankenstein protocol (Table 1). Several RNase inhibitors are commercially available, and some have been associated with reduced cDNA yields on particular single-cell platforms (*Can I use an alternative RNase inhibitor part number?*, 2026); we therefore recommend selecting one validated for the intended downstream platform.

#### Staged homogenisation improves nuclei release while limiting over-homogenisation

The Frankenstein and HCA protocols left substantial undigested tissue, whereas homogenisation in the Celegans protocol was too harsh (**Fig. 1B**). We therefore varied the number of strokes and pestle types, settling on five strokes with the loose pestle followed by three with the tight pestle, repeated until the settled suspension was clear. Nuclei were collected from the supernatant after each set, an approach adapted from the Celegans protocol (Table 1).

#### FACS is the only effective route to small-debris removal

Nematode suspensions generate considerable debris from both cuticle and lysed cells, and every additional debris-removal step costs nuclei and time. Large debris was removed most efficiently by gravity settling during dissociation followed by a single 30 µm filtration. Small debris, however, was removed effectively only by FACS: abundant cellular and cuticle debris in the pre-sorted female *H. contortus* sample (**Fig. 2B**) was largely absent after sorting (**Fig. 2C**).

#### SNIP outperforms all three existing protocols

From the same input of ten adult female *H. contortus*, SNIP recovered a mean of 31,797 ± 1,585 nuclei – 4.1-fold more than the Celegans protocol (*t*(4.0) = 18.89, *p* < 0.001), 2.3-fold more than Frankenstein (*t*(3.8) = 15.47, *p* < 0.001), and 1.7-fold more than the HCA protocol (*t*(2.8) = 5.64, *p* = 0.014), the best-performing of the three. Total RNA from the SNIP replicate (n = 1) used for extraction was 420.0 ng, equating to approximately 2.62 pg per nucleus, a 3.23-fold increase over the 0.81 pg per nucleus obtained with Frankenstein. Tapestation of RNA from nuclei isolated using the optimised protocol showed comparable RNA integrity to the existing protocols (**Fig. S1**). Sorted nuclei had smooth membranes with few showing signs of damage (**Fig. 2C**), indicating a suspension suitable for library preparation.

#### Validation of the SNIP protocol on multiple life stages of *H. contortus and T. muris*

Once optimised, we further tested our protocol with adult male and female, and third-stage (L3) larvae of *H. contortus,* and with adult female and male *T. muris* (mouse whipworm), a biologically and phylogenetically distinct parasitic nematode species (International Helminth Genomes Consortium, 2018).

Tissue disruption by scissors is not possible for microscopic stages, so for L3 larvae we substituted grinding in a dry-ice-cooled mortar and pestle. This necessarily reintroduces a transfer step between vessels, a compromise we accepted for microscopic stages, where cutting in the sample tube is not feasible. Homogenisation requirements also differed by life stage. Adult worms were fully homogenised after one set of loose-pestle strokes and three sets of tight-pestle strokes, judged by a clear supernatant. L3 larvae were more resistant and required further rounds with the tight pestle, confirmed by fluorescent staining of supernatant aliquots after each round. Additional rounds do not cost nuclei, because the supernatant is collected after each set, a feature of the protocol that makes it tolerant of variable homogenisation requirements. We found a maximum tissue volume of 20 µL per 1 mL Dounce capacity to be optimal. Importantly, additional sets of the tight Dounce tissue grinder strokes do not result in nuclei lysis because the supernatant containing nuclei is collected after each Dounce homogenisation set.

Manual nuclei counts for three independent replicates of each species and life stage tested are shown in **Fig. 3A**. For *H. contortus*, ten adult females, 20 adult males, and ∼250,000 microscopic L3 larvae were used (n = 3 biological replicates), which was approximately the same volume per tube. For *T. muris*, 25 adult female and 25 male parasites were included per replicate (n = 3 biological replicate). Although our protocol was optimised using female *H. contortus,* it proved even more effective on the female *T. muris* samples; the number of nuclei isolated per worm was significantly higher in the adult female *T. muris* (x = 26,675.33 nuclei/worm) than in *H. contortus* (x = 3,180 nuclei/worm) (*p* = 0.0001, t(4) = 15.9198). This is, in part, explained by differences in parasite size: adult female *H. contortus* worms are 20-30 mm x 300-500 µm, and adult female *T. muris* worms are 35-50 mm x 40-600 µm (**Fig. 3B**). The large range in width of *T. muris* is due to its whip-like intracellular anterior end. RNA recovery was 420.0 ng for *H. contortus* females, 37.9 ng for *H. contortus* males, 23.9 ng for *H. contortus* L3s, 901.6 ng for *T. muris* females, and 2,310.0 ng for *T. muris* males. This equates to approximately 2.62 pg/nucleus for *H. contortus* females, 0.53 pg/nucleus for *H. contortus* males, 0.17 pg/nucleus for *H. contortus* L3s, 1.66 pg/nucleus for *T. muris* females, and 2.53 pg/nucleus for *T. muris* males.

**Fig. 3.**
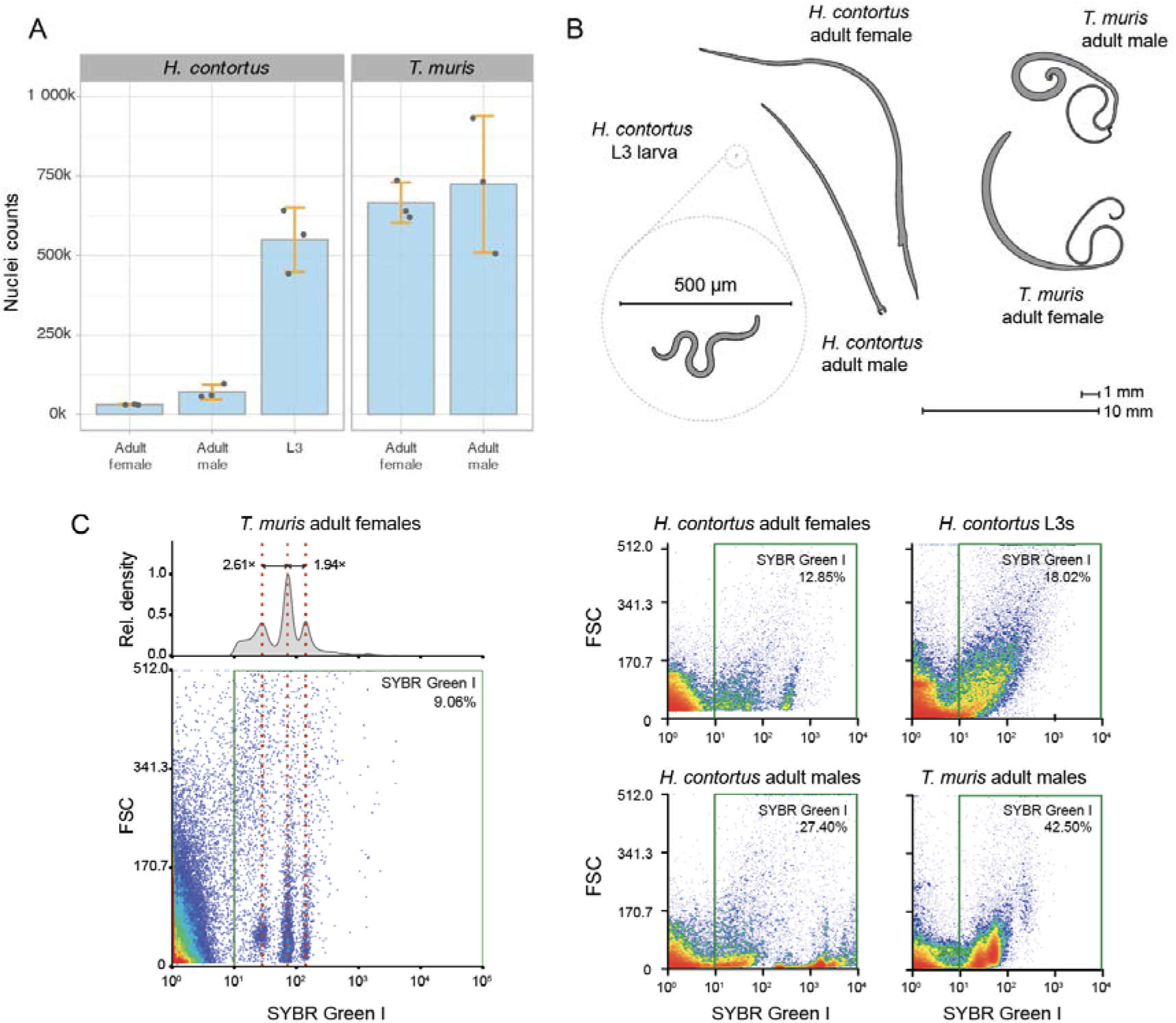
Comparison of the Single Nuclei Isolation of Parasites (SNIP) protocol on different species and life stages. A) Nuclei recovered from each species and life stage. Each replicate used ten adult female, 20 adult male or ∼250,000 L3 *H. contortus*, or 25 adult female or 25 adult male *T. muris*, matched for approximate volume per tube. Bars show the mean of three biological replicates, error bars the standard deviation, and points the individual replicates. B) Representative outlines of *H. contortus* and *T. muris* worms used in the study, highlighting the relative differences in size between life stages and species. C) Representative Fluorescence-Activated Cell Sorting (FACS) plots for each species and stage. For *T. muris* adult females, relative density (top) and FSC versus SYBR Green I fluorescence (bottom) resolve three discrete populations of increasing intensity; dotted lines mark fitted peak positions and arrows the fold change between adjacent peaks, with 95% bootstrap intervals below. Gating was determined from an unstained negative control aliquot from each sample (shown in **Fig. S3**). In all plots, the x-axis shows SYBR Green I fluorescence on a log scale (488 nm laser, 507/19 filter) and the y-axis forward scatter area (FSC-A) on a linear scale.

The FACS analysis of samples collected and fixed for snRNA-seq is shown in **Fig. 3C**. Nuclei were sorted on a SYBR Green I-positive signal (green gate), with gating determined from an unstained negative control aliquot taken from each nuclei suspension (see strategy in **Fig. S2** and per-species controls in **Fig. S3**); the proportion of total events falling within the gate ranged from 9.06% in *T. muris* adult females to 42.50% in *T. muris* adult males, reflecting substantial variation between preparations in the amount of debris carried through to sorting and underlining the need for a sorting step. Within the gate, the adult samples resolved into discrete sub-populations of differing SYBR Green I intensity, most clearly in adult female *T. muris*. Here, the SYBR Green I fluorescence of nuclei resolved into three populations of increasing intensity, with ratios between adjacent populations of 2.61× (95% bootstrap CI 2.54–2.70) and 1.94× (1.91–1.96), close to, though not precisely, the two-fold increments expected from successive rounds of genome duplication. By contrast, *H. contortus* L3 formed a single continuous distribution with no resolvable sub-populations. This pattern is consistent with differences in nuclear DNA content arising from somatic polyploidy: in *C. elegans*, intestinal and hypodermal nuclei undergo successive rounds of endoreduplication during larval development, reaching up to 32C in the adult intestine (Flemming et al., 2000; Hedgecock and White, 1985), and arrested L3 would not yet have accumulated this variation. Somatic polyploidy has also been reported in the parasitic nematode *Ascaris suum* (Anisimov et al., 1975), though it has not previously been described in *H. contortus* or *T. muris*. We note that SYBR Green I fluorescence was used here to distinguish nuclei from debris rather than to quantify DNA content, and no DNA-content standard was included; confirming the ploidy of these populations will require quantitative measurement against a calibrated reference.

#### Single-nuclei RNA library preparation and sequencing across species and life stages

To validate the SNIP protocol, we performed snRNA-seq on nuclei suspensions from both species and three life stages using the Parse Biosciences Evercode WT Mini kit, which supports up to 10,000 nuclei across 1 to 12 samples per experiment. We performed two rounds of library preparation (**Fig. S4**) and sequencing, designated “experiment 1” and “experiment 2”. **Table 2** shows the total nuclei and median genes and transcripts per nucleus for each species and life stage in both experiments, with further barcoding and sequencing metrics in **Table S1.** In experiment 1, genes and transcripts per nucleus were broadly similar across species and stages (301–446 median genes per nucleus), while total nuclei recovered varied more, highest for *T. muris* adult females (n = 1,269) and lowest for *H. contortus* adult males (n = 224). For experiment 2 we increased the number of nuclei entering barcoding, and barcoding recovery improved substantially: approximately 89,800 nuclei were loaded in experiment 1 to yield 12,750 barcoded nuclei (14%), compared with approximately 81,300 loaded and 31,000 recovered in experiment 2 (38%).

**Table 2:**
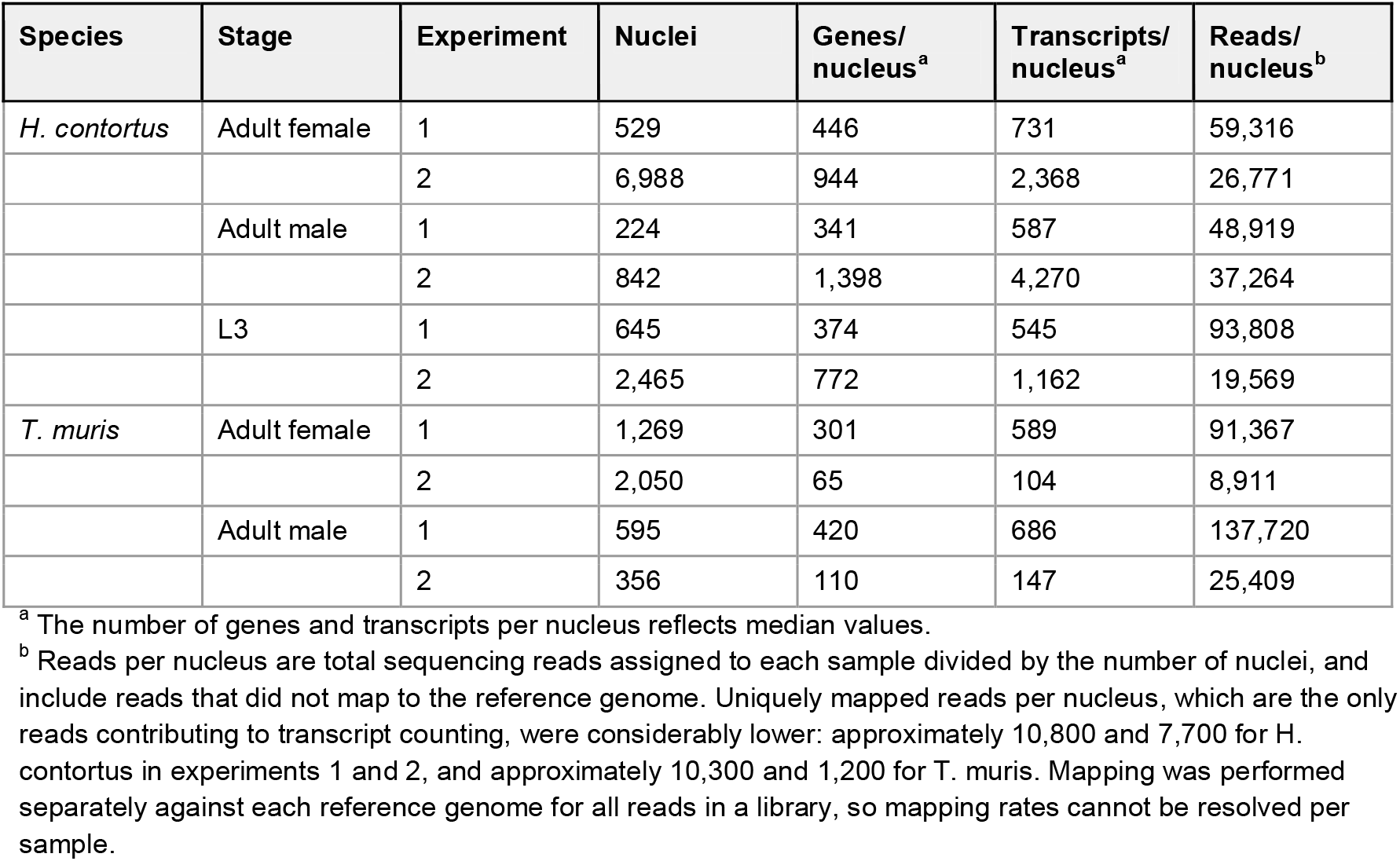
Comparison of nuclei sequencing of different species and life stages from two experiments.

Mapping efficiency, rather than sequencing depth, was the principal determinant of data recovered per nucleus.Of the reads entering alignment, 15.0% mapped uniquely to one of the two reference genomes in experiment 1 and 41.4% in experiment 2. A substantial proportion of the remainder was lost to multimapping, and this differed markedly between references: 23.5% of reads in experiment 1 and 30.3% in experiment 2 aligned to more than three loci in the *H. contortus* genome, compared with 53.5% in the *T. muris* genome in experiment 1. A further large fraction aligned to neither genome. We did not establish the origin of these reads; repetitive and ribosomal sequence, differences in assembly contiguity and annotation completeness between the two references, and host-derived material are all plausible contributors, given that both parasites were recovered from host tissue.

Because only uniquely mapped reads contribute to transcript counting, the nominal reads per nucleus in Table 2 substantially overstate the usable data. Adjusting for mapping, experiment 1 yielded approximately 10,800 uniquely mapped reads per *H. contortus* nucleus compared with approximately 7,700 in experiment 2. Despite this reduction, median genes per nucleus increased in experiment 2 for every *H. contortus* sample (adult females 446 to 944, adult males 341 to 1,398, and L3 374 to 772), indicating that differences between the two experiments reflect mapping efficiency and library complexity rather than sequencing depth. *T. muris* behaved differently: unique mapping fell to 1.4% in experiment 2, giving approximately 1,200 uniquely mapped reads per nucleus and correspondingly few genes (65 and 110 for adult females and males). Barcode and cDNA quality scores were also lower for this library (Q30 0.79–0.83, compared with 0.95–0.97 in experiment 1), suggesting a library or sequencing quality issue rather than a difference in the nuclei themselves.

To test whether the differences between experiments affected downstream analysis, we clustered *H. contortus* nuclei from both experiments together (**Fig. S5**). Genes per nucleus were consistently high across all clusters except Cluster 0, which was dominated by experiment 2 L3 nuclei (**Fig. S5A–B**). All samples contributed nuclei to every cluster (**Figs. S45,C–G**), although the proportion contributed to each varied, as expected, given that the experiments were not designed to capture all cell types and the number of nuclei sequenced per sample was small relative to a comprehensive atlas. Overall, we saw no clear difference in cluster structure between the two experiments.

#### Single-nuclei RNA sequencing identifies distinct cellular populations in different parasite life stages

The sequencing was designed to validate the library preparation approach across a range of sample inputs rather than to generate a comprehensive atlas, which would require substantially more nuclei per sample. Even at this scale, analysis of transcript expression per species resolved clusters of nuclei by life stage and sex (**Fig. 4A** for *H. contortus* and **Fig. 4F** for *T. muris*), and clusters that were shared or unique across life stages, consistent with cell- and tissue-specific variation (**Fig. 4B,G**).

**Fig. 4:**
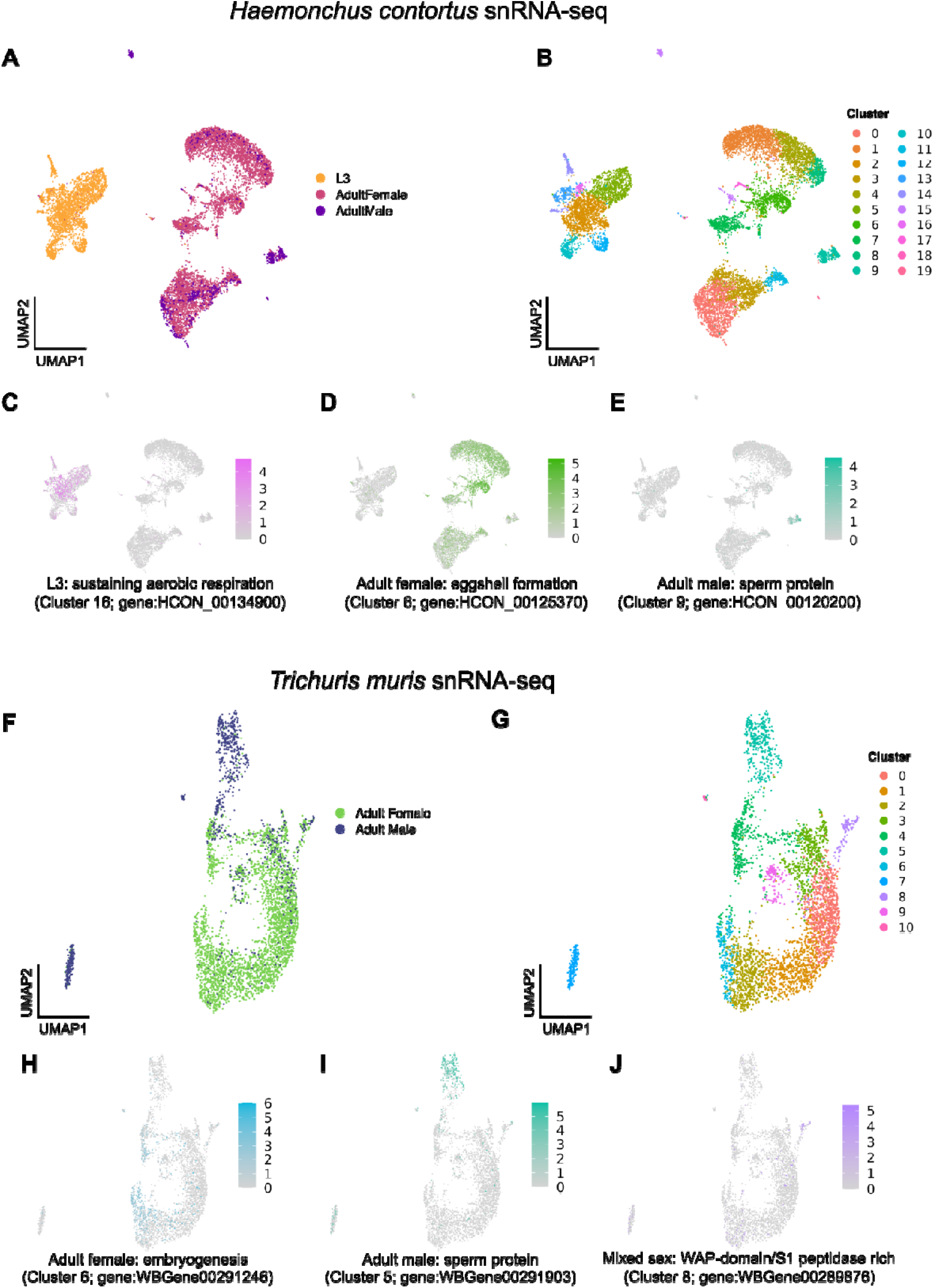
Single-nuclei RNA sequencing of different life stages of *Haemonchus contortus* and *Trichuris muris.* Uniform Manifold Approximation and Projections (UMAPs) of snRNA-seq data from (A-E) *H. contortus* and (F-J) *T. muris* coloured by (A, F) life stage or nuclei cluster (B, G). In total, 20 nuclei clusters were predicted from three life stages (adult male, adult female, L3) of *H. contortus*, and 11 nuclei clusters from two life stages (adult male, adult female) of *T. muris*. In C-E and H-J, we highlight three examples of biologically meaningful clusters from both datasets, demonstrating sex- and stage-specific gene expression (colour scale: log-normalised expression counts) and inferred functional roles based on *Caenorhabditis elegans* orthology.

We inferred the putative function of each cluster from gene set enrichment and orthology to the model free-living nematode *C. elegans*. This revealed stage- and tissue-specific gene expression, including those putatively involved in aerobic respiration required outside of the host in *H. contortus* L3 (**Fig. 4C**), and eggshell formation (**Fig. 4D**) and sperm (**Fig. 4E**) specific to adult females and males, respectively. Similar stage- and tissue-specific expression was observed in *T. muris*, with clusters provisionally associated with gametogenesis in females (**Fig. 4H**) and sperm proteins in males (**Fig. 4I**), and a mixed-sex cluster enriched for WAP-domain/S1 peptidase expression (**Fig. 4J**).

These assignments should be interpreted cautiously, for two reasons. First, the substantial evolutionary distance between *C. elegans* and the species examined here means that marker genes used to characterise a cell or tissue type in one may not reliably identify the same type in the other (Wong et al., 2024). Second, the assignments rest on relatively sparse data per nucleus: as described above, a large proportion of reads did not map to the reference genomes, and median transcript counts per nucleus were correspondingly low, so clusters are defined by a limited number of detected genes. The identities proposed here are therefore hypotheses to be tested rather than established cell-type assignments.

Considering the sparse functional annotation information for non-model species such as parasitic nematodes (Doyle, 2022), data such as these represent both a challenge and an opportunity: improving genomic resources for these species is a prerequisite for building the comprehensive cell and nuclei atlases needed to understand their biology.

## Discussion

In this study, we evaluated three existing protocols for isolating single nuclei for snRNA-seq and optimised a method for recovering nuclei from parasitic nematodes (SNIP). Our motivation for this work was to create robust cell atlases that will enhance our understanding of parasitic nematode development, illuminate the tissues and organs that mediate interactions with their hosts, and identify novel drug targets and mechanisms of drug resistance. Here, we discuss the rationale for our approach, the challenges faced at both the bench and in data analysis, and the ongoing limitations to be overcome to apply snRNA-seq to parasitic nematodes specifically and, more generally, to apply cellular-resolution technologies to other complex and enigmatic non-model organisms.

The use of single-cell and single-nuclei RNA sequencing to understand the fundamental biology of organisms at cellular resolution is advancing rapidly. Although there are successful examples of cell atlases for specific life stages or tissues of multicellular parasites using scRNA-seq (Attenborough et al., 2024; Diaz Soria et al., 2024, 2020; Henthorn et al., 2023; Li et al., 2021; Pollo et al., 2024; Puckelwaldt et al., 2024; Tyagi et al., 2024), we specifically chose to use snRNA-seq for several reasons. First, we aim to collect and analyse parasites at different life stages over time and in response to perturbations, such as drug treatment exposure. Whereas scRNA-seq typically requires rapid processing of fresh tissue and live cells from recovery through to library preparation, snRNA-seq enables sample preservation via freezing, allowing collection of samples over time that can be processed together, thus reducing library and sequencing batch effects. Here, we demonstrated the versatility of the protocol by sampling multiple species and life stages, each derived from different hosts (*H. contortus* in sheep and *T. muris* in mice) and using different sampling strategies (invasive and non-invasive sampling from hosts). Moreover, freezing parasites enabled sample collection and shipment from geographically distant collaborators, rather than requiring physical presence with the specimens at the time of collection for immediate processing. Such an approach opens the possibility of sampling other non-model organisms from diverse or difficult-to-access locations worldwide, provided they can be frozen shortly after collection.

The second reason for a nuclei-focused approach is the challenge of tissue dissociation into living cells. Both scRNA-seq and snRNA-seq aim to mirror the transcriptome of cells in tissue, and thus, the recovery of “healthy” cells is paramount. However, stress caused by a combination of physical and/or enzymatic treatment, temperature, and time can alter transcriptional profiles and result in artefacts during data analysis (van den Brink et al., 2017; Wu et al., 2019). Therefore, effective cellular dissociation of an organism or tissue for scRNA-seq requires a balance between sufficient effort (physical or enzymatic) to dissociate and sufficient care not to damage cells or nuclei so they can be processed. The dissociation of frozen tissue to collect nuclei, in which transcriptional profiles are less susceptible to change and therefore better reflect the transcriptome signature of the cells, can be preferred for difficult-to-dissociate tissues that require greater physical forces to be applied during dissociation. We anticipated this to be important for dissociating parasitic nematodes, many of which are protected by a strong acellular cuticle and sheath and are very resistant to disruption. Furthermore, our optimised protocol improved the dissociation time; the original protocols tested took ∼40-45 minutes to complete, whereas the optimised protocol took ∼30 minutes.

Other advantages of snRNA-seq over scRNA-seq include the capture of a greater diversity of cell types (Slyper et al., 2020), likely due to differential susceptibility to damage between cell types, and the physical constraints of cell size of commonly used microfluidic technologies, for example, 10x Genomics has a recommended maximum cell size limit of 30 µm and a maximum physical microfluidic channel width of 50-60 µm. Cell sizes are known to vary considerably in nematodes; for example, during development of the model free-living nematode *C. elegans*, the intestine increases substantially in tissue volume through to adulthood, primarily because intestinal cells grow in size and become multinucleated and polyploid rather than through a large increase in cell number (Hedgecock and White, 1985). We anticipated similar variation in the closely related *H. contortus*, and in specialised tissues such as the elongated stichosome of *T. muris* (O’Sullivan et al., 2021). Consistent with this, FACS profiles of adult *H. contortus* and *T. muris* nuclei resolved into discrete sub-populations of differing SYBR Green I intensity that were absent from arrested L3 larvae (**Fig. 3C**), as would be expected if nuclei of differing DNA content accumulate during development. Moreover, approximately 30% of *C. elegans* somatic nuclei are contained in large, multinucleated cells due to cellular fusion during development to form complex tissues (Podbilewicz, 2006). Due to their unusual cell morphology properties, these cell types are likely to be better represented in snRNA-seq data than in scRNA-seq data.

Our choice of the Parse Biosciences Evercode platform, based on Split Pool Ligation-based Transcriptome sequencing (SPLiT-seq) (Rosenberg et al., 2018), followed from the same reasoning. Unlike droplet-based platforms such as 10x Genomics Chromium, combinatorial barcoding allowed nuclei preparations from individual life stages to be collected at different times, fixed, and stored frozen before library preparation, so that samples collected weeks apart could be processed together. In our experiments, storage before library preparation ranged from 1 to 106 days without apparent loss of performance, and the manufacturer specifies stability for up to six months (*Evercode Fixation and WT v2 User Guides*, 2023).

This removes a substantial source of technical variation between samples and replicates, which is particularly valuable when working across life stages that cannot be obtained simultaneously. We also considered the capacity to pool up to 384 samples in a single experiment when scaling up, the reduced ambient RNA relative to droplet-based approaches, the use of both poly(A) and random primers to capture mRNA and non-coding RNAs (Rosenberg et al., 2018), and the absence of any requirement for specialised equipment. Parse is not the only route to fixing and storing samples before processing; Roux et al. (Roux et al., 2023), for example, used methanol-fixed *C. elegans* cells with 10x Chromium Single Cell 3′ chemistry.

We overcame several challenges during protocol development. For adult stages of parasitic worms, we found that chopping frozen tissue in a single tube on dry ice resulted in more efficient dissociation and lysis than thawing and mincing tissue. However, dissociating microscopic *H. contortus* L3 larvae, an environmentally robust free-living life stage with a sheath surrounding its external cuticle (Cantacessi et al., 2010), proved more challenging. We explored several approaches, including a *C. elegans* nuclei isolation protocol by Cao *et al*. (Cao et al., 2017) that used shearing forces from pipetting samples through a 21 G needle for initial tissue disruption. However, in our hands, the *H. contortus* L3 larvae did not shear, whereas the larger adult stages persistently clogged the needle. Our approach of pre-grinding frozen L3 with a porcelain mortar and pestle, followed by a combination of loose and tight Dounce strokes with recovery of nuclei between sets of strokes, substantially improved nuclei recovery. Throughout the protocol, nuclei were lost at each step. We tackled this by recovering nuclei during homogenisation in stages, which prevented excessive mechanical damage and thus maximised the collection of intact nuclei. Once collected, nuclei needed to undergo filtration to remove large and small debris; we found that a single wash step with a 30 µm filter and a filter wash removes sufficient small and large debris to prevent clogging of the FACS machine capillaries while minimising loss of nuclei. The Celegans protocol recommended using Flowmi Cell Strainers, which attach directly to a pipette; however, we found they were easily clogged by the amount of debris in parasitic nematode nuclei suspensions and could not be easily washed to improve nuclei recovery.

Furthermore, while FACS removes small debris, reveals diverse nuclei populations, and provides another count measurement, it can also lead to loss of nuclei and decreased nuclei quality (Alfieri et al., 2022). The value of FACS as a filtration step depends on the degree of debris present, and where debris is minimal it may be worth comparing sequencing outcomes with and without a sorting step. We also observed nuclei loss during fixation using the Parse Biosciences Evercode protocol for snRNA-seq. Parse Biosciences recommends that high nuclei numbers be fixed to account for loss (*Evercode Fixation and WT v2 User Guides*, 2023); however, the loss seems sample-dependent and fixation requires optimisation to minimise its impact. For example, the *T. muris* adult male nuclei suspension showed a 69.33% loss of nuclei, compared with only 11.88% for *T. muris* adult female samples. Because we omitted the two 40 µm straining steps specified in the fixation protocol in order to maximise retention, this loss cannot be attributed to filtration. This result suggests that some nuclei are more fragile or may indicate more extensive nuclear blebbing in the *T. muris* male nuclei, causing an already compromised nuclear envelope to lyse during fixation (Srivastava et al., 2021). Male *T. muris* nuclei also yielded more RNA per nucleus than females (2.53 versus 1.66 pg), although total RNA largely reflects the greater number of nuclei recovered from males. Whether higher RNA content per nucleus and greater fragility during fixation share a common basis is not resolved by these data, particularly as RNA was quantified before fixation. Similarly, the Parse Biosciences WT Mini library preparation requires high sample input (∼55,000 cells/nuclei) at specific concentrations (between 298 and 3,575 cells/nuclei per µL, depending on the number of samples), which can be challenging to achieve with low-biomass samples. One potential solution is to combine and concentrate fixed nuclei samples to obtain the proper input concentration, with the caveat that it can result in substantial overall nuclei loss due to additional centrifugation and resuspension.

Losses were not confined to the bench. A substantial proportion of sequencing capacity was lost before transcript counting, with relatively low rates of uniquely mapped reads to either reference genome, with a further large fraction either multimapping or failing to align at all. Nuclei yield is therefore not the only constraint on applying snRNA-seq to non-model organisms. Reference contiguity and annotation completeness, repetitive and ribosomal sequence, and host-derived material in parasites recovered from host tissue all reduce the usable data obtained per unit of sequencing, and the magnitude of this loss is unlikely to be apparent without inspecting mapping statistics directly. For organisms recovered from a host, and for species whose genomes remain fragmented or incompletely annotated, we therefore recommend assessing mapping rate on a pilot library before committing to deep sequencing.

We acknowledge several limitations to our study. While we have gained insights from the two parasitic nematode species tested, nuclei isolation from other species, and extension to other non-model organisms, may present distinct challenges and require species-specific optimisation, particularly regarding the number of Dounce tissue grinder strokes. Some variation in nuclei counts may reflect differences in the susceptibility of particular tissues and cell types to disruption; until denser atlases are available and cellular and nuclear proportions are better understood, this is difficult to address. Where individual or small numbers of samples can be processed without freezing and thawing after fixation, higher nuclei counts and more input material for library preparation will likely be achieved. RNA yields were measured on a single replicate per protocol and per sample, and are therefore indicative rather than statistically supported. The sub-populations observed by FACS are consistent with differences in nuclear DNA content, but SYBR Green I was used here to distinguish nuclei from debris rather than to quantify DNA content, and confirming ploidy will require measurement against a calibrated standard. Finally, the cluster annotations presented here are provisional: they rest on orthology to *C. elegans* and on gene set enrichment, neither of which substitutes for direct validation of cell-type identity in these species.

## Conclusions

We have built on existing approaches to develop an optimised protocol for single nuclei isolation of parasitic nematodes for snRNA-seq. Validation across several diverse species and multiple life stages demonstrates versatility, and a critical evaluation of steps provides a substantial head start for using this protocol to isolate nuclei from other non-model organisms. The increasing accessibility of cellular-resolution techniques and optimised protocols will advance the understanding of organisms, such as parasitic nematodes, from fundamental biology to infection and disease, and toward novel approaches to control them.

## Code and data availability

Reference genomes and annotations used in the analysis can be accessed from WormBase ParaSite (https://parasite.wormbase.org/). Raw sequencing data can be accessed from the European Nucleotide Archive (ENA) under the study accession ERP140984. The code used to analyse sequencing data can be accessed from https://github.com/stephenrdoyle/single_cell_optimisation. Any additional information required to reanalyse the data reported in this paper is available from one of the lead contacts.

## Contributions

Conceptualisation: SKB, SRD

Investigation: SKB, AM, ACB, MRK, BLN, KM, EH, SRD

Methodology: SKB

Data Curation: SKB

Formal analysis: SKB, SRD

Resources: DJB, MAD-C

Formal Analysis: SKB, SRD

Funding Acquisition: SRD, DJB, MAD-C

Project administration: SKB, DJB, MAD-C, SRD

Supervision: SRD

Writing – Original Draft Preparation: SKB, SRD

Writing – Review & Editing: All authors

## Acknowledgements

We gratefully acknowledge the technical support and advice of Ian Pittam from Parse Biosciences, the Pathogen Informatics group at the Wellcome Sanger Institute for informatics support, and the Helminth Genomics group at the Wellcome Sanger Institute for constructive feedback on the analyses and manuscript. This work is supported by a UKRI Future Leaders Fellowship to SRD [MR/T020733/1] and the Wellcome Trust (UK) through core funding to the Wellcome Sanger Institute (UK) [206194]. AAM and DJB receive underpinning national capacity funding from the Rural and Environment Science and Analytical Services (RESAS) division of the Scottish Government. MAD-C was supported by the Sir Henry Dale Fellowship jointly funded by the Wellcome Trust and the Royal Society [222546/Z/21/Z]; the Wellcome Trust [203151/Z/16/Z, 203151/A/16/Z] and the UKRI Medical Research Council [MC_PC_17230]. For the purpose of Open Access, the author has applied a CC BY public copyright licence to any Author Accepted Manuscript version arising from this submission.

