## Supplementary Information for "Optimisation of single-nuclei isolation and RNA sequencing of parasitic nematodes"

Figure S1. TapeStation analysis of RNA extracted from nuclei derived from established and optimised protocols.	2

Representative Agilent HS RNA TapeStation gel of RNA extracted from the nuclei suspension of each protocol, as shown in Fig. 1D. Here, the TapeStation gel of RNA extracted from the optimised protocol (SNIP) is shown for comparison.	2

Figure S2. Fluorescence-Activated Cell Sorting (FACS) gating strategy for nuclei.	3

Figure S3. FACS plots of sample-matched unstained negative controls.	4

Figure S4. TapeStation HS D5000 fragment trace analysis of Parse sub-libraries and final libraries.	5

Figure S5. Extended analysis of Haemonchus contortus experiments	6

Table S1. Additional nuclei barcoding and sequencing metrics from the two Parse Evercode WT mini kit experiments.	7

Table S2. Mapping rates to each genome from experiments 1 and 2	8

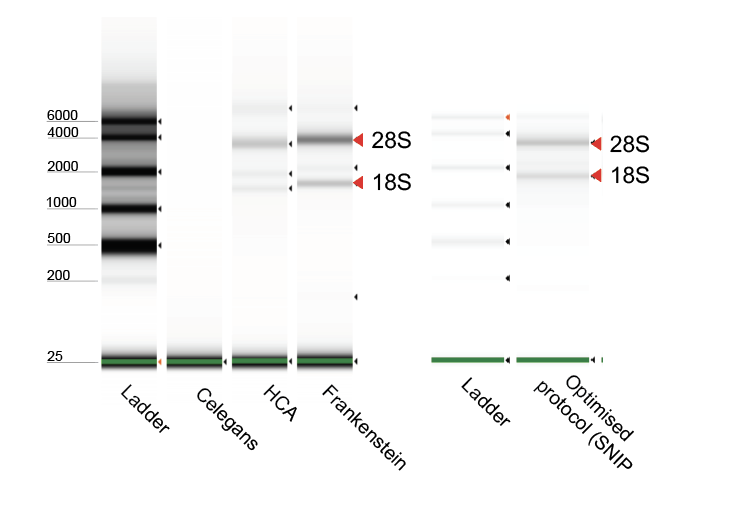

### Figure S1. TapeStation analysis of RNA extracted from nuclei derived from established and optimised protocols.

### Representative Agilent HS RNA TapeStation gel of RNA extracted from the nuclei suspension of each protocol, as shown in Fig. 1D. Here, the TapeStation gel of RNA extracted from the optimised protocol (SNIP) is shown for comparison.

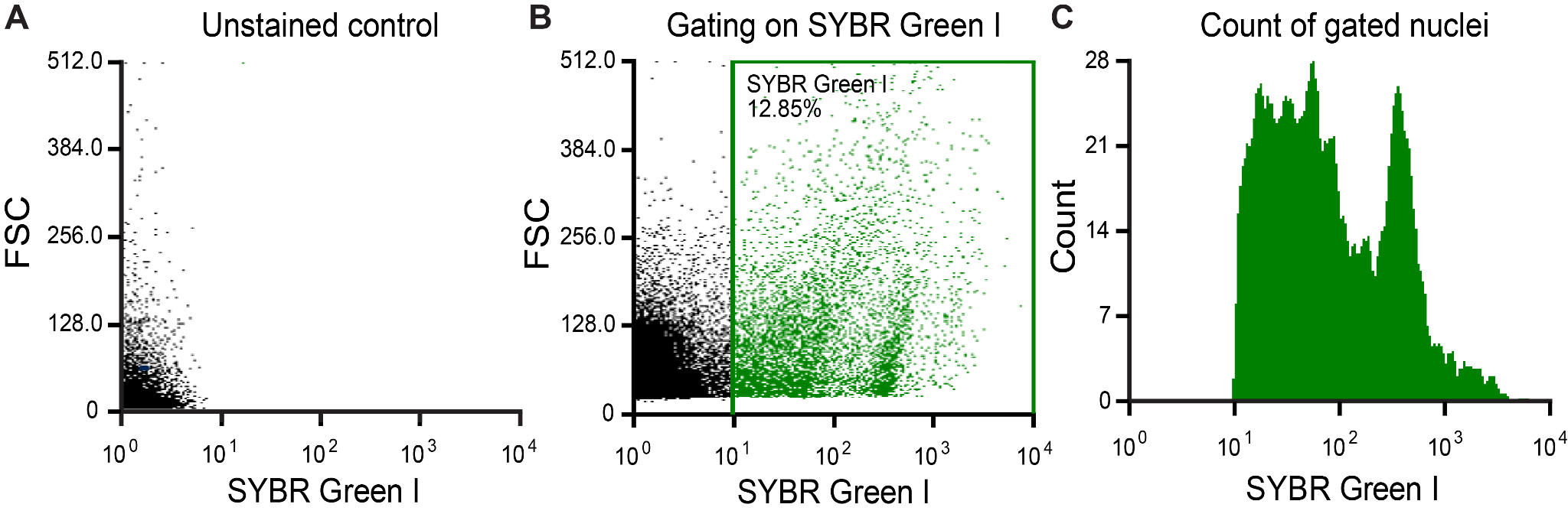

### Figure S2. Fluorescence-Activated Cell Sorting (FACS) gating strategy for nuclei.

Small debris from the nuclei suspensions was removed using FACS. A) A 50 µL aliquot of each nuclei suspension was retained unstained as a negative control to establish fluorescence detection thresholds. B) The remaining suspension (~500 µL) was stained with 5 µL 1,000× SYBR Green I (ThermoFisher) to a final concentration of approximately 10×, and nuclei were gated on a positive fluorescent signal relative to the unstained control from the same suspension (gate shown in green). Cellular and cuticle debris remain unstained and fall outside the gate. The x-axis shows SYBR Green I fluorescence on a log scale (488 nm laser, 507/19 filter); the y-axis shows forward scatter area (FSC-A) on a linear scale.

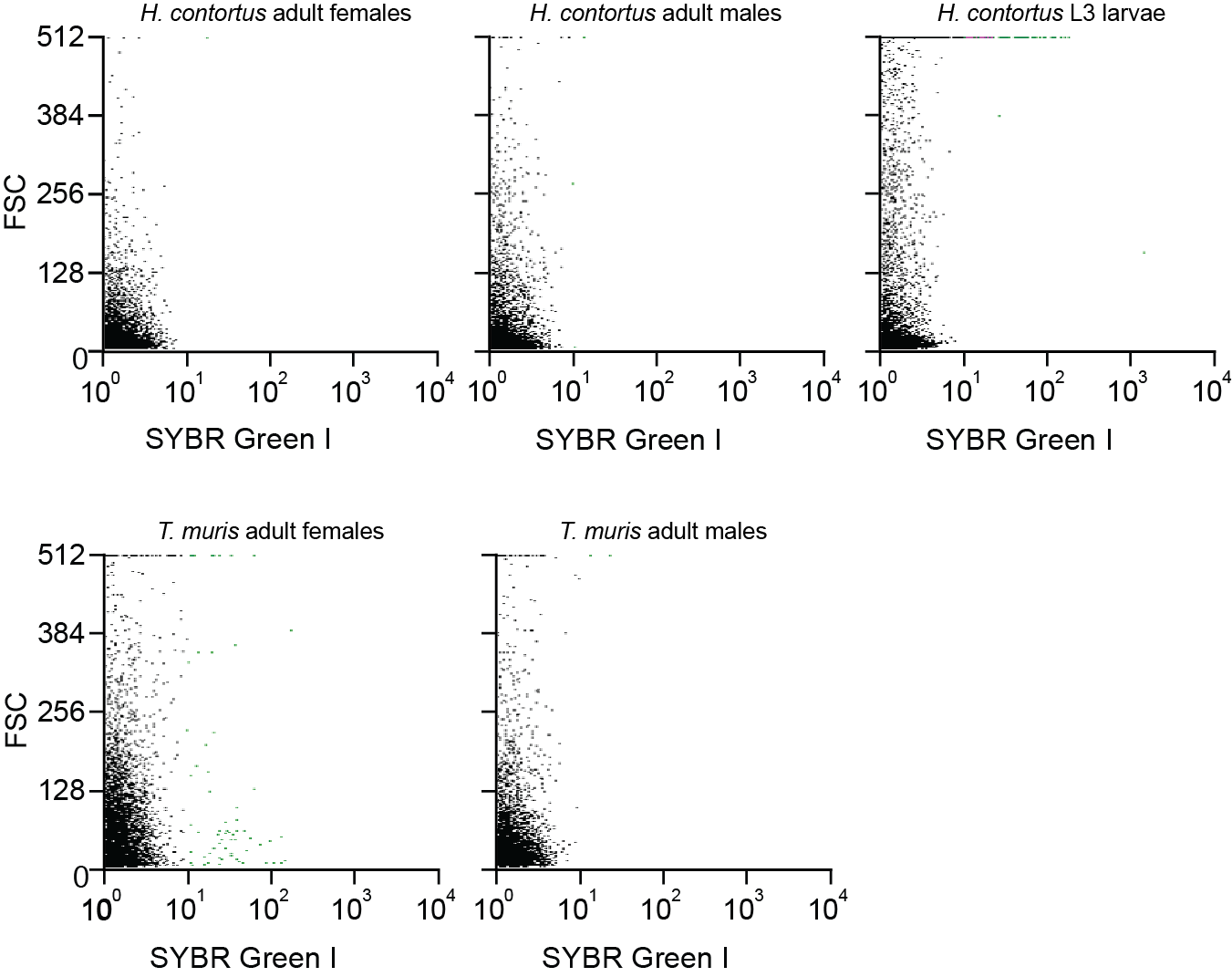

### Figure S3. FACS plots of sample-matched unstained negative controls.

Unstained aliquots from each of the five nuclei preparations - *H. contortus* adult females, adult males and L3, and *T. muris* adult females and adult males - isolated using the SNIP protocol. Each was used to set the SYBR Green I-positive gate for its corresponding stained sample in Figure 3C. Axes as in Figure S1.

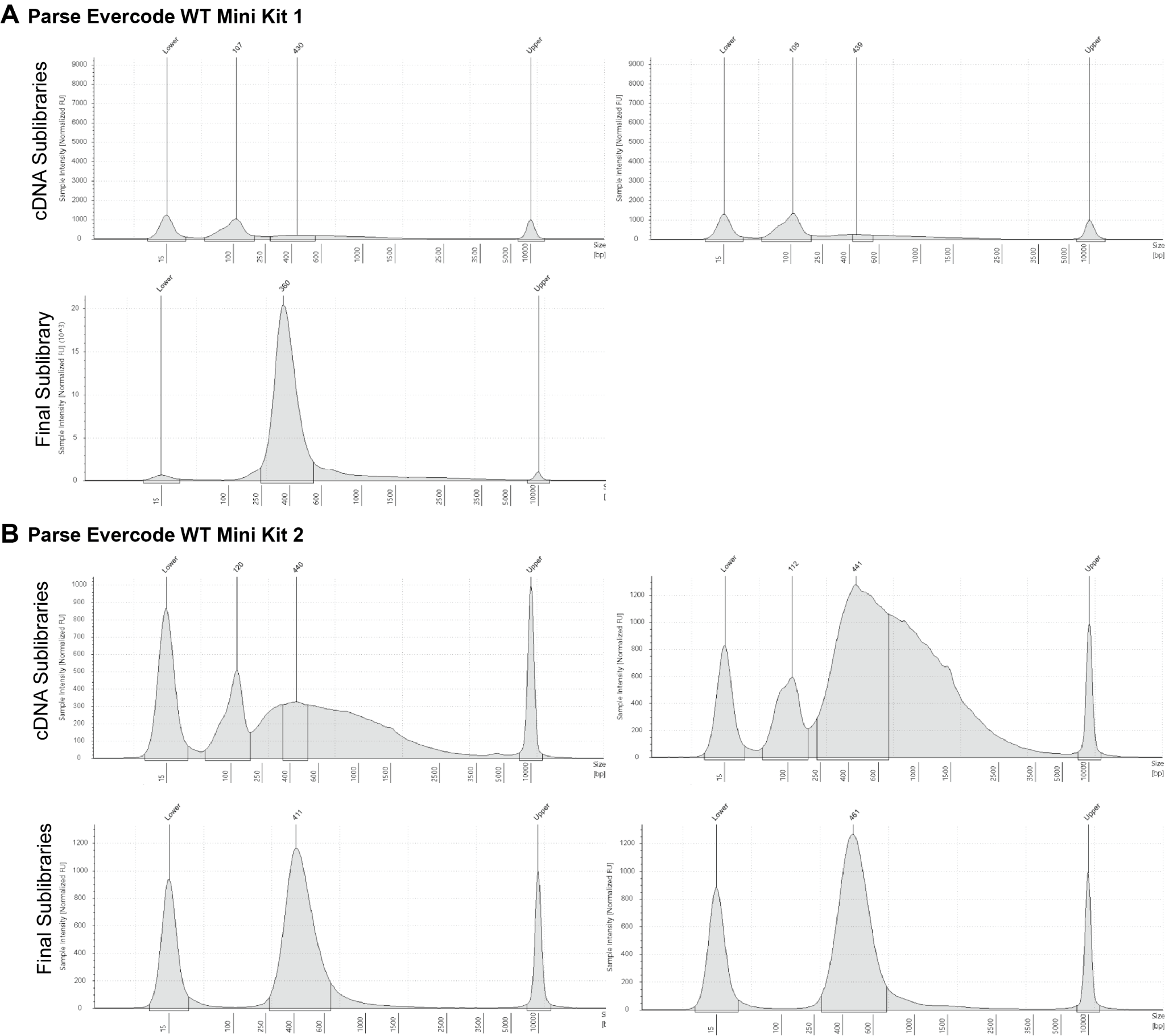

### Figure S4. TapeStation HS D5000 fragment trace analysis of Parse sub-libraries and final libraries.

A) Parse Evercode WT Mini Kit run 1 fragment traces of the two cDNA sub-libraries and the resulting final sub-library from the combined sub-libraries. Qubit measurements for sub-libraries 1 and 2 were 2.22 ng/µl and 2.12 ng/µl, respectively. We combined the sub-libraries, and the remaining 60.76 ng cDNA was used for fragmentation, end repair, and A-tailing. The final library was quantified using a Qubit to 18.6 ng/µl. B) Parse Evercode WT Mini Kit run 2 fragment traces of the two cDNA sub-libraries in a 1:2 dilution. Qubit measurements for sub-libraries 1 and 2 were 6 ng/µl and 12 ng/µl, respectively. 100 ng of each sub-library was used for fragmentation, end repair, and A-tailing. The two final indexed sub-libraries were assessed on the TapeStation in a 1:5 dilution. Qubit measurements for final libraries 1 and 2 were 6.34 ng/µl and 9.54 ng/µl, respectively.

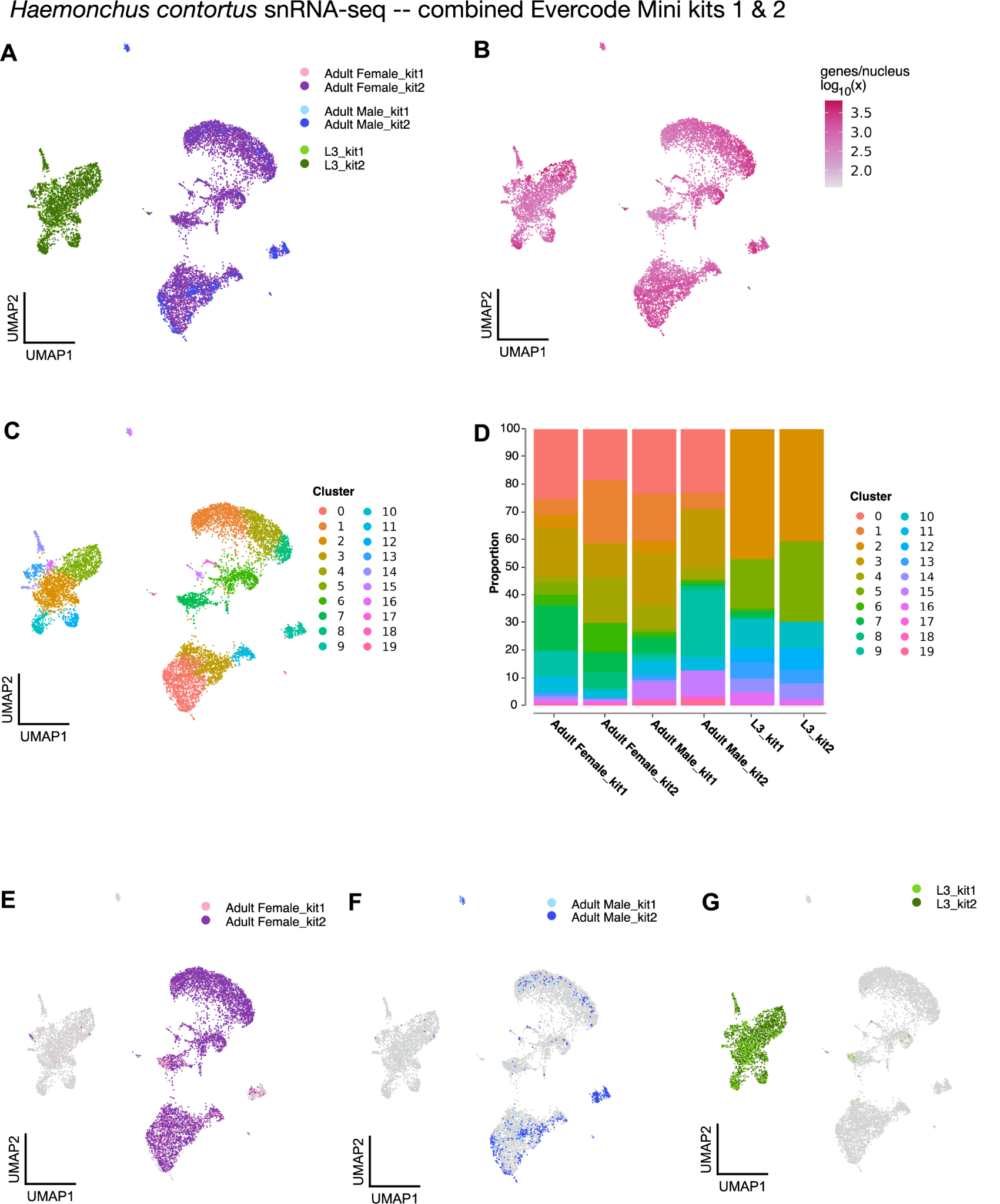

### Figure S5. Extended analysis of *Haemonchus contortus* experiments

To explore differences between Parse Evercode WT Mini Kit 1 and 2, we combined nuclei sequenced across all *H. contortus* samples from both experiments using Harmony. A UMAP coloured by sample is shown in A, with a heatmap of transcripts per nucleus shown in B. Overall cluster classifications are presented in C, and the proportion of nuclei contributing to each cluster is shown in D. Sample-specific UMAPs, separated by experiment, are displayed in E-G.

### Table S1. Additional nuclei barcoding and sequencing metrics from the two Parse Evercode WT mini kit experiments.

| **Species** | **Stage** | **Exp** | **Post-thaw count^a^** | **Days frozen** | **Nuclei added to barcoding** | **Barcoded nuclei^c^** | **Nuclei sequenced** | **Genes/ nucleus** | **Transcripts/ nucleus** | **Reads/ nucleus** | **Sequencing saturation** |
| --- | --- | --- | --- | --- | --- | --- | --- | --- | --- | --- | --- |
| *H. contortus* | Adult female | 1 | 95,550 | 68 | 16,380 | 12,750 | 529 | 446 | 731 | 59,316 | 0.668 |
|  |  | 2 | 53,500 | 106 + 11^b^ | 18,360 | 31,000 | 6,988 | 944 | 2,368 | 26,771 | 0.216 |
|  | Adult male | 1 | 118,650 | 8 | 18,367 | 12,750 | 224 | 341 | 587 | 48,919 | 0.699 |
|  |  | 2 | 43,750 | 1 + 11^b^ | 7,875 | 31,000 | 842 | 1,398 | 4,270 | 37,264 | 0.210 |
|  | L3 | 1 | 330,575 | 8 | 18,361 | 12,750 | 645 | 374 | 545 | 93,808 | 0.712 |
|  |  | 2 | 68,000 | 1 + 11^b^ | 18,360 | 31,000 | 2,465 | 772 | 1,162 | 19,569 | 0.212 |
| *T. muris* | Adult female | 1 | 278,775 | 36 | 18,367 | 12,750 | 1,269 | 301 | 589 | 91,367 | 0.710 |
|  |  | 2 | 32,250 | 7 | 18,362 | 31,000 | 2,050 | 65 | 104 | 8,911 | 0.234 |
|  | Adult male | 1 | 119,175 | 67 | 18,367 | 12,750 | 595 | 420 | 686 | 137,720 | 0.725 |
|  |  | 2 | 72,000 | 7 | 18,365 | 31,000 | 356 | 110 | 147 | 25,409 | 0.230 |

^a^ Post-thaw nuclei counting was performed using a CytoFLEX for experiment 1 and manually with a disposable haemocytometer for experiment 2.

^b^ Two replicates isolated on separate days were combined to achieve the necessary concentration and nuclei starting material.

^c^ Barcoded nuclei reflects a pooled count per experiment.

Nuclei sequenced is the number of nuclei called by split-pipe v1.9.0 (default cell-calling cutoff). Genes/nucleus and transcripts/nucleus are medians; reads/nucleus is the mean number of reads allocated per nucleus (all reads in a sublibrary, including those without a valid barcode, apportioned to samples by their wells’ valid-barcode reads). Experiment 1 is a single sublibrary and Experiment 2 combines two sublibraries.

### Table S2. Mapping rates to each genome from experiments 1 and 2

| **Experiment** | **Reference genome** | **Reads entering alignment** | **Uniquely mapped** | **Multimapped (≤3 loci)** | **Mapped to >3 loci** |
| --- | --- | --- | --- | --- | --- |
| 1 | *H. contortus* | 230,067,012 | 15,155,424 (6.59%) | 2,157,002 (0.94%) | 54,019,929 (23.48%) |
| 1 | *T. muris* | 230,067,012 | 19,231,879 (8.36%) | 6,594,436 (2.87%) | 123,091,451 (53.50%) |
| 2 | *H. contortus* | 198,787,250 | 79,362,492 (39.92%) | 9,204,883 (4.63%) | 60,261,802 (30.31%) |
| 2 | *T. muris* | 198,787,250 | 2,872,171 (1.44%) | 925,557 (0.47%) | 12,507,035 (6.29%) |

Nuclei from both species were barcoded together within each experiment, so we mapped all reads against each reference genome in turn; percentages therefore reflect the total reads entering alignment, not reads originating from that species. Only uniquely mapped reads (MAPQ 255) contribute to transcript counting. Reads mapping to more than three loci were discarded by STAR (--outFilterMultimapNmax 3).
